# Treatment with anti-inflammatory viral serpin modulates immuno-thrombotic responses and improves outcomes in SARS-CoV-2 infected mice

**DOI:** 10.1101/2022.09.09.507363

**Authors:** Liqiang Zhang, Yize (Henry) Li, Karen Kibler, Simona Kraberger, Arvind Varsani, Julie Turk, Nora Elmadbouly, Emily Aliskevich, Laurel Spaccarelli, Bereket Estifanos, Junior Enow, Isabela Rivabem Zanetti, Nicholas Saldevar, Efrem Lim, Kyle Browder, Anjali Wilson, Fernando Arcos Juan, Aubrey Pinteric, Aman Garg, Savanah Gisriel, Bertram Jacobs, Timothy L. Karr, Esther Borges Florsheim, Vivek Kumar, John Wallen, Masmudur Rahman, Grant McFadden, Brenda G. Hogue, Alexandra R. Lucas

## Abstract

1.

Severe acute respiratory distress syndrome (ARDS) during SARS-CoV-2 (severe acute respiratory syndrome coronavirus-2) infection, manifests as uncontrolled lung inflammation and systemic thrombosis with high mortality. Anti-viral drugs and monoclonal antibodies can reduce COVID-19 severity if administered in the early viremic phase, but treatments for later stage immuno-thrombotic syndrome and long COVID are limited. *Ser*ine *p*rotease *in*hibitors (SERPINS) regulate activated proteases during thrombotic, thrombolytic and immune responses. The myxoma poxvirus-derived Serp-1 protein is a secreted immunomodulatory serpin that targets activated coagulation and complement protease pathways as part of a self-defense strategy to combat viral clearance by the innate immune system. When purified and utilized as an anti-immune therapeutic, Serp-1 is effective as an anti-inflammatory drug in multiple animal models of inflammatory lung disease and vasculitis. Here, we describe systemic treatment with purified PEGylated Serp-1 (PEGSerp-1) as a therapy for immuno-thrombotic complications during ARDS. Treatment with PEGSerp-1 in two distinct mouse-adapted SARS-CoV-2 models in C57Bl/6 and BALB/c mice reduced lung and heart inflammation, with improved clinical outcomes. PEGSerp-1 significantly reduced M1 macrophage invasion in the lung and heart by modifying urokinase-type plasminogen activator receptor (uPAR) and complement membrane attack complex (MAC). Sequential changes in urokinase-type plasminogen activator receptor (uPAR) and serpin gene expression were observed in lung and heart with PEGSerp-1 treatment. PEGSerp-1 is a highly effective immune-modulator with therapeutic potential for treatment of severe viral ARDS with additional potential to reduce late SARS-CoV-2 complications related to immune-thrombotic events that persist during long COVID.

**Significance:** Severe acute respiratory distress syndrome (ARDS) in SARS-CoV-2 infection manifests as uncontrolled tissue inflammation and systemic thrombosis with high mortality. Anti-viral drugs and monoclonal antibodies reduce COVID-19 severity if administered early, but treatments for later stage immuno-thrombosis are limited. *Ser*ine *p*rotease *in*hibitors (SERPINS) regulate thrombotic, thrombolytic and complement pathways. We investigate here systemic treatment with purified poxvirus-derived PEGSerp-1 as a therapeutic for immuno-thrombotic complications in viral ARDS. PEGSerp-1 treatment in two mouse-adapted SARS-CoV-2 models (C57Bl/6 and BALB/c) significantly reduced lung and heart inflammation and improved clinical outcomes, with sequential changes in thrombolytic (uPAR) and complement expression. PEGSerp-1 is a highly effective immune-modulator with therapeutic potential for immune-thrombotic complications in severe viral ARDS and has potential benefit for long COVID.

## 2. Introduction

Excessive clotting, bleeding and inflammation increase lung damage, hypoxia and mortality in the severe Acute Respiratory Distress Syndrome (ARDS) associated with the later stages of Coronavirus-2 (SARS-CoV-2) lung infections. Mortality rates up to 20-45% have been reported for hypoxic patients admitted to intensive care unit (ICU) (1–4). With severe virus infections, ARDS induces aggressive and uncontrolled inflammation that fails to shut off, thereby causing further damage with lung consolidation, hypoxia, widespread coagulopathy and increased mortality (1–9). In healthy physiologic states, there is a balance between reciprocal activation of the thrombotic and thrombolytic cascades and reciprocal interactions with immune responses, providing the homeostasis necessary to regulate and prevent excessive or uncontrolled coagulation and immune response activation (6,9–11). ARDS with severe viral lung infections can induce excess and damaging immune responses and coagulopathies, with a loss of this normal homeostasis. As with SARS-CoV-2, other respiratory viruses like SARS-CoV-1, Middle East Respiratory Syndrome (MERS) (5) and Influenza (10–22) can all induce damaging imbalance of the immune and thrombotic/coagulation responses.

Current anti-viral therapies can be highly effective post-exposure treatments for SARS-CoV-2 infections if administered in the early stages of viremia, reducing the risk of ICU admission, but the later stage immuno-thrombotic complications of SARS-CoV-2 infections remain difficult to treat with 40-50% mortality in severe cases (1–4). Corticosteroids provide benefit in patients with hypoxia and more advanced lung disease but are only partially effective and are not recommended in early stages of COVID-19 (1–4,23,24). Anticoagulants such as heparin have had variable benefit and there can be marked resistance in SARS-CoV-2 intensive care unit (ICU) admissions (25–28). Targeted monoclonal antibodies against individual pro-inflammatory cytokines can also have limited therapeutic benefits. There are also long-term sequelae, termed long COVID, which may be associated with the failure to turn off these immuno-thrombotic complications in multiple organs, not just the lungs. There is at present no prevention or treatment for long COVID (28–31).

Thrombolytic and complement pathways are comprised of serine protease cascades that govern coagulation and innate immune responses. Increased activation levels of multiple serine protease cascades are detectable in severe SARS-CoV-2 lung infections, serving as markers for higher risk of respiratory distress, systemic thrombosis, long term clinical sequelae and increased mortality (1–5, 31–34). Soluble urokinase-type plasminogen activator receptor (suPAR) and activated complement are two clinical markers for risk of progression to ARDS and ICU admission in SARS-CoV-2 infections. uPA and tissue-type plasminogen activator (tPA) are two commonly measured circulatory thrombolytic serine proteases (the clot dissolving cascade) that help regulate overall coagulation pathways. The uPA/ uPAR complex is positioned at the leading edge of migratory leukocytes that mediate the innate cellular immune responses and is a central mediator in inflammation, immunity, and coagulation. Additionally, uPA/uPAR activates plasmin and also matrix metalloproteinases, thereby increasing inflammatory macrophage invasion as well as growth factor activation. tPA is the primary thrombolytic serine protease and increased soluble tPA is also reported in severe SARS-CoV-2 infections (31–35). tPA operates by binding and activating plasminogen activated receptors, PAR (30,31) The complement cascade is also comprised of multiple activated serine proteases with a sequential activation designed to form the C5b-9 membrane attack complex (MAC), an effector for innate and acquired immune cell responses (37). There is also a mutual feedback interaction between the complement and thrombotic pathways that can cause runaway coagulopathies if left unchecked.

The uPA, tPA, thrombin and complement cascades are all regulated by mammalian serpins, representing 2% or more of circulating plasma proteins. Serpins have been examined both as biomarkers for disease as well as potential therapeutics for genetic disorders, serpinopathies and for infection-related coagulopathies. The mammalian serpin, plasminogen activator inhibitor-1 (PAI-1, SERPINE1) binds uPA and tPA and circulating PAI-1 levels are elevated, together with tPA, in SARS-CoV-2 infected patients (36). Anti-thrombin III (SERPINC1) has been investigated as a treatment for severe bacterial sepsis and disseminated intravascular coagulation (38). ATIII targets the thrombotic cascade and, while producing a trend toward benefit, did not achieve significance in treatment for bacterial sepsis. Other clinical trials are in progress examining the mammalian serpins, alpha1 antitrypsin (A1AT, SERPINA1) (39) and Complement-1 inhibitor (C1-Inh, SERPING1) (40) and as treatments for severe SARS-CoV-2, both with a range of inhibitory functions.

Many large DNA viruses have evolved highly potent immunomodulatory proteins over eons of selective evolutionary pressures, typically as a self-protective strategy against clearance by activated innate immune cells of various classes, thus providing a natural reservoir for biologics designed to block key pivotal regulatory pathways in the immune response (41,42). We have investigated select members of these highly effective virus-derived immune-modulating proteins as a source for new antiinflammatory therapeutics designed to treat the damaging inflammation and coagulopathies in severe viral infections and to investigate molecular mechanisms driving severe ARDS (43). Serp-1 is a myxoma virus-derived secreted 55kDa serpin glycoprotein that operates in virus-infected tissues by protecting the virus from attack by activated myeloid cells (44). Myxoma is a rabbit poxvirus with high mortality in European rabbits, but non-pathogenic for humans. Purified Serp-1 protein, however, acts as a species-nonspecific inhibitor of systemic inflammation by binding and inhibiting multiple activated serine proteases in the coagulation and complement pathways (45). Serpins bind activated serine proteases, specifically targeting cellular and extracellular sites of serine protease enzyme activation. Systemic treatment with purified Serp-1 downregulates trafficking of migratory macrophages and lymphocytes into damaged tissues via binding to uPAR (46,47). uPAR (CD87) is a glycosylphosphatidylinositol (GPI)-linked surface protein that sits in a large lipid raft of proteins that interact with integrins, chemokine receptors, vitronectin, growth factors, low density lipoprotein receptor-related protein (LRP) and complement receptors on the cell surface, altering cell activation and motility (48). Serp-1 also binds and inhibits several complement proteases (49). Bolus systemic treatment with clinical grade Serp-1 has proven safe and effective in a Phase IIA randomized dose escalating clinical trial, performed at 7 sites in the US and Canada, in unstable coronary syndrome patients with stent implants (50). In this trial, Serp-1 administered intravenously together with current standard of care of acute unstable coronary syndromes, significantly reduced early markers for heart damage, troponin (TN) and creatinine kinase MB (CK-MB), with minimal adverse events at the therapeutic dose.

A modified PEGylated serpin, PEGSerp-1 has been recently developed with improved halflife and activity (49). This modified Serp-1 still effectively binds uPA, tPA and thrombin and a series of complement proteases, specifically C1(q,r and s), factor B, C3 and C4 proteases as assessed by mass spectrometry. PEGSerp-1 represents a new class of immune modulating virus-derived biologic with extensive proven efficacy in a wide spectrum of systemic inflammatory diseases, including gamma herpesviral vasculitis (43) and lung infection, pristane-induced diffuse alveolar hemorrhage (DAH)(49), numerous animal models of vascular injury and allograft transplants (43). In both the virus infection and DAH models, serpin treatment altered expression levels of numerous coagulation and complement pathway proteases and serpins (43,49).

In this report, we assess the efficacy of systemic PEGSerp-1 treatment to target immune-thrombotic pathways and treat severe SARS-CoV-2 viral lung infections in two mouse-adapted models, MA10 in BALB/c mice (51) and MA30 in C57BL/6 mice (52).

## 3. Results

### PEGSerp-1 treatment of MA10-infected BALB/c mice

As a preliminary screen, PEGSerp-1 treatment was assessed in SARS-CoV-2 MA10 infected BALB/c mice (N = 16, Table 1)(51). In this pilot study, PEGSerp-1 was given daily with early follow up at 48 hours. Lung consolidation was compared in uninfected and in SARS-CoV-2 MA10 infected mice, with and without PEGSerp-1 treatments (Figure 1, Panel 1). PEGSerp-1 was given daily with early follow up at 48 hours. In this initial study, PEGSerp-1 treatment produced a trend toward a reduction in lung consolidation and inflammation as detected on histological analysis, but which did not reach 0.05 significance level (Figure 1, Panel 1 A-C; *p* = 0.1073). PEGSerp-1 treated mice with SARS-CoV-2 MA10 infection had similar levels of consolidation as seen in mice without viral infection (Fig 1, Panel 1C). On immunohistochemical analysis (IHC), both detectable iNOS positive M1 macrophage counts (Figure 1, Panel 1D-F; *p* < 0.05) and F4/80 positive macrophage counts (Figure 1, Panel 1G; *p* < 0.012) were significantly reduced in the lung by the PEGSerp-1 treatment. Lung sections from uninfected controls with saline or PEGSerp-1 treatments had minimal areas of iNOS positive macrophage cell counts (Figure 1, Panel 1F). PEGSerp-1 did not produce adverse effects in uninfected or infected mice.

**Figure 1.**
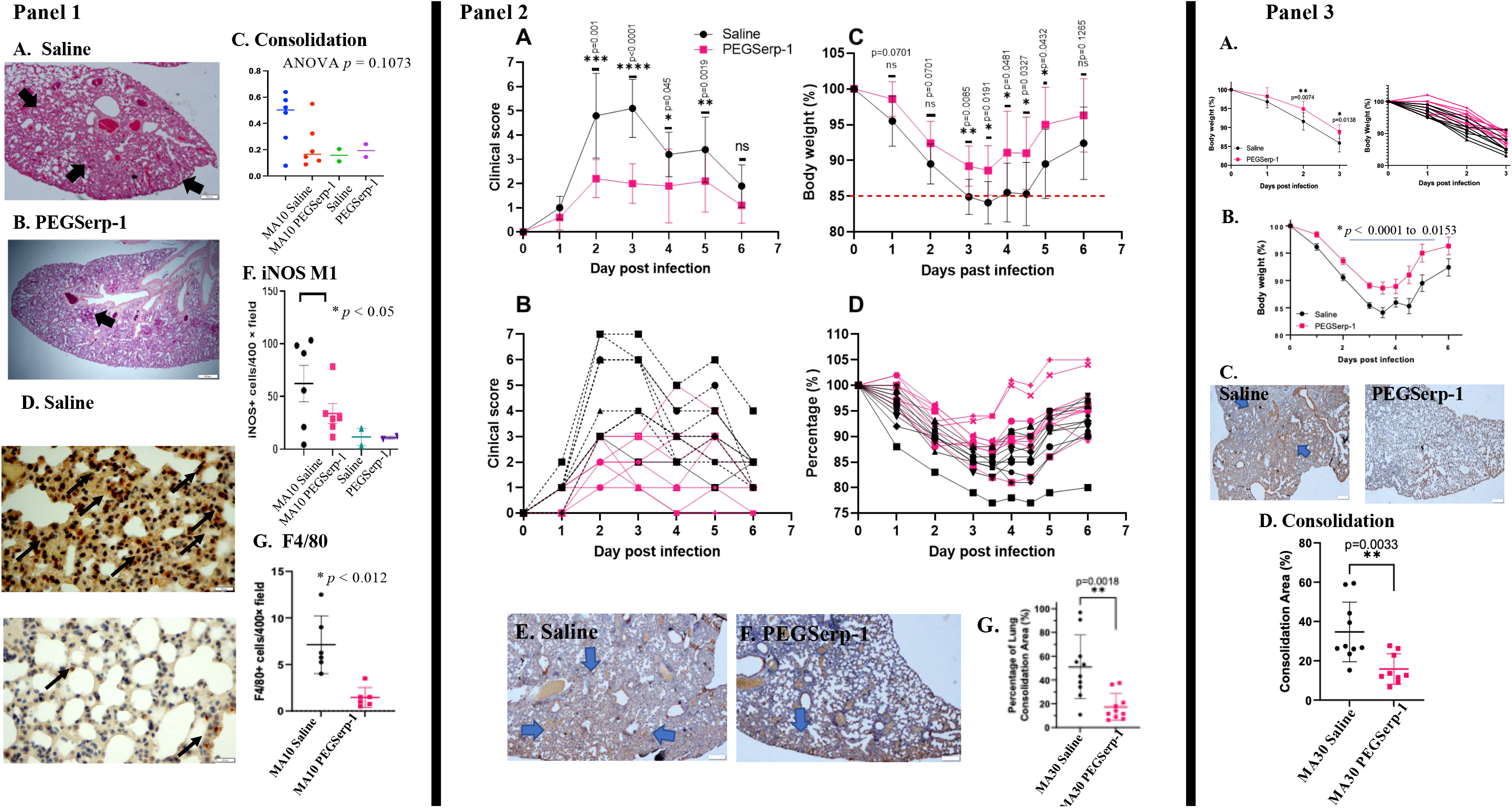
PEGylated Serp-1 treatment improves clinical score and weight gain and reduces lung consolidation in SARS-CoV-2 MA10 infected BALB/c and MA30 infected C57Bl/6 mouse models. ***Panel 1*** –Pilot study, SARS-CoV-2 MA10 infected BABLB/c mice at 48 hours follow up. PEGSerp-1 treatment reduced lung consolidation and iNOS + macrophage infiltration. H & E stained histology sections of lung illustrating consolidation in Saline treated mice (A) reduced by PEGSerp-1 treatment (B). Graph of measured consolidation area divided by total lung area (C; p = 0.1073). iNOS M1 stained macrophage invasion in Saline control treated mice (D) is significantly reduced with PEGSerp-1 treatment (E) Graphs illustrate significant reductions in mean iNOS positive M1 macrophage (F; *p* < 0.05).and F4/80 macrophage (G; p < 0.012) positive stained cell counts. Mean for positively stained cell counts in 3 HPF per mouse. ***Panel 2*** – SARSCoV-2 MA30 infection in C57BL/6 mice 7 days follow up. PEGSerp-1 treatment significantly improved mean (A) and individual (B) clinical scores (; *p* < 0.045 to 0.0001, days 2-5) as well as Mean (C) and individual weights (D)(*p* < 0.0481- 0.0085 days 3-5). Micrographs illustrate lung consolidation in Saline (E) and PEGSerp-1 (F) treated lungs. Graph of (G) mean consolidation area divided by total area at 7 days follow up. (G) (Red squares – PEGSerp-1, Black circles - Saline) ***Panel 3***. SARS-CoV-2 MA30 infection in C57Bl/6 mice 4 days follow up. PEGSerp-1 treatment significantly improved weight gain during the second 4 day follow up study. Improved mean (A) and individual (B) weights with PEGSerp-1 treatments in MA30 infected mice *(p* < 0.138- 0.0074), Combined analysis of mice in 4 day and 7 day follow up indicate consistent improved weight gain with PEGSerp-1 treatment (B). Micrographs illustrate lung consolidation in Saline and PEGSerp-1 treated (C) MA30 infected mice. Consolidation is significantly reduced at 4 days follow up with PEGSerp-1 (D; *p* < 0.0033- < 0.001). Large blue arrows indicate areas of consolidation in H&E stained lung sections (Mag 2X); Small black arrows point to brown iNOS positive macrophage on IHC stained lung sections (Mag 40X).

**Table 1.**
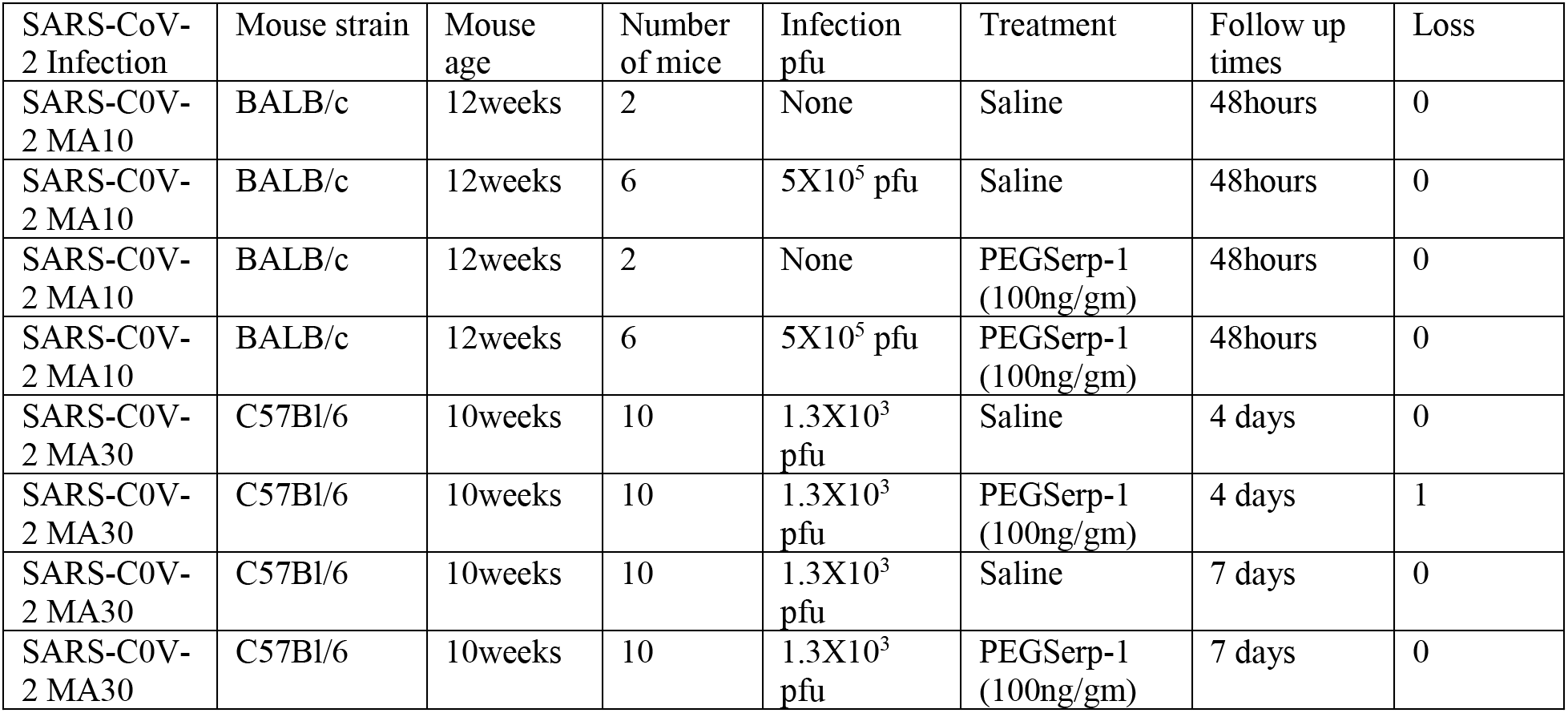
SARS-CoV-2 Mouse Infections Models.

| SARS-CoV-2 Infection | Mouse strain | Mouse age | Number of mice | Infection pfu | Treatment | Follow up times | Loss |
| --- | --- | --- | --- | --- | --- | --- | --- |
| SARS-C0V-2 MA10 | BALB/c | 12weeks | 2 | None | Saline | 48hours | 0 |
| SARS-C0V-2 MA10 | BALB/c | 12weeks | 6 | 5X10<sup>5</sup> pfu | Saline | 48hours | 0 |
| SARS-C0V-2 MA10 | BALB/c | 12weeks | 2 | None | PEGSerp-1 (100ng/gm) | 48hours | 0 |
| SARS-C0V-2 MA10 | BALB/c | 12weeks | 6 | 5X10<sup>5</sup> pfu | PEGSerp-1 (100ng/gm) | 48hours | 0 |
| SARS-C0V-2 MA30 | C57Bl/6 | 10weeks | 10 | 1.3X10<sup>3</sup> pfu | Saline | 4 days | 0 |
| SARS-C0V-2 MA30 | C57Bl/6 | 10weeks | 10 | 1.3X10<sup>3</sup> pfu | PEGSerp-1 (100ng/gm) | 4 days | 1 |
| SARS-C0V-2 MA30 | C57Bl/6 | 10weeks | 10 | 1.3X10<sup>3</sup> pfu | Saline | 7 days | 0 |
| SARS-C0V-2 MA30 | C57Bl/6 | 10weeks | 10 | 1.3X10<sup>3</sup> pfu | PEGSerp-1 (100ng/gm) | 7 days | 0 |

The SARS-CoV-2 MA10 model in BALB/c mice produced lung consolidation with histological evidence for lung infection, but did not produce significant weight loss or clinical symptoms (scores) in our preliminary study, though previous studies exhibited a more pathogenic phenotype in BALB/c mice (51). To further assess the potential efficacy of PEGSerp-1 for treating SARS-CoV-2, PEGSerp-1 treatment was further examined in a second mouse adapted SARS-CoV-2 model, the MA30 infection model in C57Bl/6 mice (52,53).

### PEGSerp-1 Treatment of MA30-infected C57Bl/6 mice

PEGSerp-1 treatment was next assessed in SARS-CoV-2 MA30 infected C57Bl/6 mice using a Lethal dose 50 (LD50) inoculation calculated at 1.3-1.4X10^3^pfu (YL,BH). MA30-infected mice were treated with daily intraperitoneal injections (IP) of either PEGSerp-1 or control saline and followed for 7 days (N = 20 mice, Table 1). The SARS-CoV-2 MA30 infection model produced a significant systemic response in saline control treated C57Bl/6 mice with increased clinical score and weight loss in infected, mice (Figure 1, Panel 2). MA30-infected mice treated with saline control displayed significant clinical symptoms with respiratory distress (Figure 1, Panel 2A, B) and weight loss (Figure 1, Panel 2C, D) on days 2 to 5 post infection followed by gradual improvement on day 6. The clinical score was a combined scoring for weight loss together with scores for hunching, ruffling/sneezing, and labored breathing. There was a significant reduction in the clinical score in PEGSerp-1 treated SARS-CoV-2 MA30-infected mice on days 2-5 (Figure 1, Panel 2A, B; with a range of statistical scores from *p* < 0.045 – 0.001), together with significantly reduced weight loss on days 3-5 with PEGSerp-1 treatments (Figure 1, Panel 2C,D; *p* < 0.0432 – 0.0084). Overall efficacy for PEGSerp-1 treatment was greatest at 3-5 days after infection. Lung consolidation was significantly reduced with PEGSerp-1 treatments (Figure 1, Panel 2E-G) as measured by mean area of lung consolidation divided by total lung area in stained sections, calculated for multiple reads and sections per mouse (Figure 1, Panel 2G; *p* < 0.0018). An outside blinded SARS lung pathology score was also performed and indicated similar trends toward reduced lung disease with PEGSerp-1 treatment *(p* = 0.085; Supplemental Figure 1).

A third cohort of mice was assessed at 4 days follow up, with and without PEGSerp-1 treatment (N = 20), to provide a second, independent analysis of PEGSerp-1 treatment during SARS-CoV-2 MA30 infection and also to provide an earlier timepoint for lung pathology assessment (Figure 1, Panel 3). PEGSerp-1 treatment again reduced weight loss on days 2 and 3 after infection (Figure 1, Panel 3A; *p* < 0.0074 - 0.0138). To assess for variability in the model, analysis of combined data from both 4 day and 7 day follow up studies demonstrated preserved significance, further supporting a consistent benefit with PEGSerp-1 treatment (Figure 1, Panel 3B). Reduced weight loss for the day 4 follow up group with PEGSerp-1 treatment was again associated with a significant reduction in lung consolidation on histopathological assessment, corroborating the prior 7 day follow up study (Figure 1, Panel 3C, D). No mice died during the 7 days follow up. One mouse died immediately upon intranasal (IN) inoculation of SARS-CoV-2 in this second group due to an anesthesia complication prior to receiving treatment.

The results overall from the two cohorts of MA30 infection mice indicate consistent, significantly improved weight gain and clinical scores at days 2-4 for PEGSerp-1 treatment of infected mice with follow up at days 4 and 7 post-infection. Lung consolidation was also significantly reduced with measured reductions in areas of lung consolidation at both days follow up.

### Analysis of inflammatory cell invasion in SARS-CoV-2 lung infections

Cellular immune responses were also examined in both MA10 and MA30 SARS-CoV-2 infection models by immunhistochemical (IHC) analysis to measure neutrophil, macrophage and T cell infiltrates in lung sections from both models. iNOS-positive M1 macrophage cell counts were significantly reduced in the MA10 SARS-CoV-2 infected mice treated with PEGSerp-1 at 48 hours follow up as noted (Figure 1, Panel 1D-G; p < 0.05). iNOS positive cell counts were also significantly reduced in SARS-CoV-2 MA30 infected/treated mice at day 4 (Figure 2, Panel 1A, B; *p* < 0.045) and day 7 (Figure 2, Panel 1 C, D; *p* < 0.0052) follow up as measured on lung micrograph sections, with a greater reduction detected at 7 days. Arg1-positive M2 macrophage cell counts were not altered on day 4 (Figure 2, Panel 1D; *p* = 0.2921), but were significantly increased on day 7 (Figure 2, Panel 1E; *p* < 0.0414) in MA30 infected mice with PEGSerp-1 treatment, consistent with a progressive anti-inflammatory effect (Figure 2, Panel 1E). Neutrophil counts, measured by the Ly6G marker, had a non-significant trend toward a decrease in PEGSerp-1 treated infected mice on day 4 (Supplemental Figure 1, Panel 1A, *p* = 0.086), but no change on day 7 (Figure 1, Panel 1B; *p* = 0.6570). CD3 positive T cell staining was not altered on day 4 (Supplemental Figure 1, Panel1C; *p* = 0.1058). CD4 staining was not altered on day 4 with PEGSerp-1 treatment (Figure 2, Panel 1G; *p* = 0.5013). In contrast, CD4 positively stained T cells were significantly reduced by PEGSerp-1 treatment on day 7 in MA30 infected mice (Figure 2, Panel 1H, *p* < 0.0454). There was a nonsignificant trend toward an increase in CD8 positive T cells at day 4 (Figure 2, Panel 1I; *p* = 0.1503), but at 7 days follow up CD8 T cells were significantly reduced in PEGSerp-1 treated infected mice, similar to trends observed with CD4 stained cells (Figure 2, Panel 1J; *p* < 0.0303).

**Figure 2.**
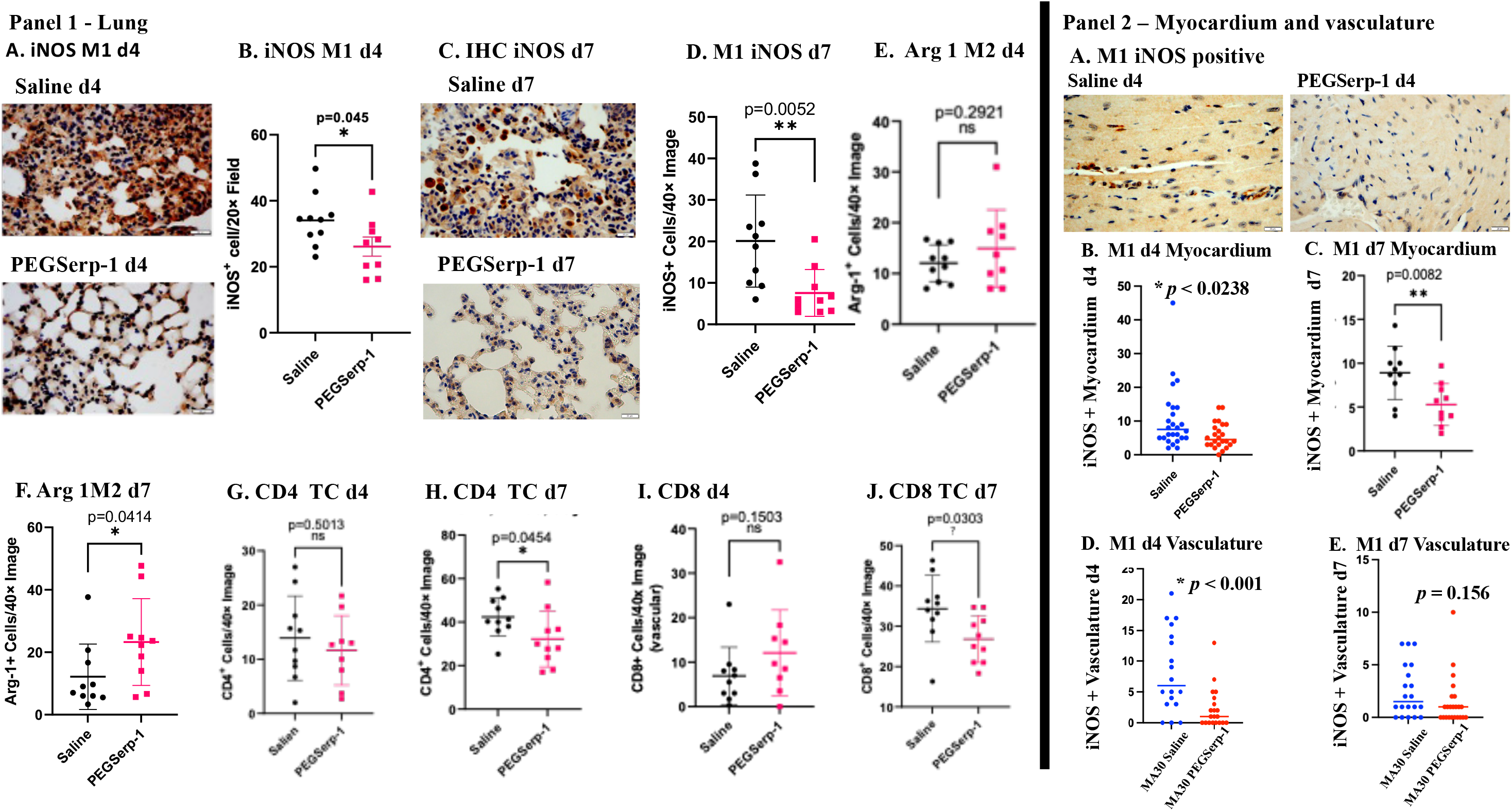
PEGSerp-1 treatment significantly reduced immune cell infiltrates in in SARS-CoV-2 MA30 infected C57BL/6 mouse models at 4 and 7 days follow up. ***Panel 1*** –iNOS positive M1 macrophage cell infiltrates on IHC stained lung sections in Saline and PEGSerp-1 (A) treated mice with significantly reduced IHC positive cell counts at 4 days follow up (B; *p* < 0.045). iNOS positive infiltrates in IHC stained lung sections with Saline or PEGSerp-1 (C) treatment with significantly reduced IHC positive cell counts at 7 days follow up (D; *p* < 0.0052). Arginase1 positive M2 macrophage IHC cell counts with a trend toward an increase at 4 days (E; *p* = 0.2921) and a significant increase at 7 days (F; *p* < 0.0414) with PEGSerp-1 treatment. CD4 + T cell counts are not reduced at 4 days (G; *p* =0.5031) but are reduced at 7 days (H; *p* < 0.0454) with PEGSerp-1 treatments. CD8 positive T cell counts on IHC stained sections have a trend toward an increase at 4 days (I; *p* = 0.1503) and a significant reduction at 7 days (J; *p* < 0.0303) with PEG Serp-1 treatments. ***Panel 2*** - Myocardium and vascular staining detect reduced overall iNOS positive M1 macrophage cell counts on IHC micrographs of myocardium in PEGSerp-treated SARS-CoV-2 MA30 infected C57BL/6 mice. iNOS positive cells in Saline and PEGSerp-1 treated mice at 4 days follow up (A), Graphs illustrate significantly reduced mean iNOS positive cell counts in the myocardium at 4 days (B) and 7 days (C). iNOS positive perivascular cell counts are significantly reduced at 4 days (D; *p* < 0.001), but not 7 days follow up (E; *p* = 0.156). IHC stained sections 40X). Graphs illustrate mean ± SD for positively stained cells counts on IHC stained sections (Black Saline, Red – PEGSerp-1)..

### Parallel assessment of myocardial and vascular inflammation in MA10- and MA30-SARS-CoV-2 infected mice demonstrated reduced inflammatory cell invasion with PEGSerp-1 treatments

The entire cardiovascular system has been associated with widespread inflammatory immune cell responses and micro-thrombotic vascular occlusions during SARS-CoV-2 infections in humans. We examined the effects of SARS-CoV-2 MA30 infections in hearts isolated from virus-infected C57Bl/6 mice, with and without PEGSerp-1 treatments, at days 4 and 7 post-infection follow up. Inflammation was detected in the myocardium and in associated vasculature but was much less prominent than in the lungs from Saline treated control SARS-CoV-2 infected mice (Figure 2, Panel 2). Inflammatory cell infiltrates were detectable in the myocardium in isolated pockets often in the pericardium or leaflets and in perivascular spaces.

PEGSerp-1 significantly reduced detectable iNOS-positive M1 macrophage cell counts in the myocardium at 4 (Figure 2, Panel 2A-C; *p* < 0.0238) and 7 days (Figure 2, Panel 2D;*p* < 0.0082) follow up. In the small vessels in the myocardium, iNOS-positive M1 macrophage counts were significantly decreased at day 4 (Figure 2, Panel 2E,*p* < 0.001), and on day 7 (Figure 2, Panel 2F,*p* = 0.156). Nonspecific CD3 positive T cell counts were not significantly reduced by PEGSerp-1 at day 7 in the myocardium (Supplemental Figure 1, Panel 2E; *p* = 0.113) and in the vasculature (Supplemental Figure 1, Panel 2G, *p*= 0.5482) of virus-infected mice. CD4+ T cell counts were also not significantly altered by PEGSerp-1 in the myocardium or vasculature (Supplemental Figure 1, Panel 2F, H; *p* = 0.6408 and 0.9404)). Ly6G positive cell counts were also not significantly changed by PEGSerp-1 in the myocardium or vasculature.

### Immunohistochemical analysis for uPAR and complement

As a serpin, PEGSerp1-binds to the uPA/uPAR complex (Supplemental Figure 2)(44,45,47–49) and also a wide range of complement cascade proteases, as demonstrated on Mass spectrometry pull down in a hemorrhagic lung model (49)(Table 2). uPAR is commonly upregulated and expressed on the leading edge of migratory leukocytes and can also be cleaved and released into the blood as soluble uPAR (suPAR) (46,47). Thus, increased uPAR expression at the cell surface may be reflected in elevated cleaved uPAR, as soluble uPAR (suPAR). Changes in detectable uPAR and C5b-9 membrane attack complex (MAC) levels were measured by IHC analysis of lung sections from SARS-CoV-2 infected mice treated with PEGSerp-1.

Cell associated uPAR staining was significantly reduced at day 2 follow up in SARS-CoV-2 MA10-infected BALB/c mice treated with PEGSerp-1 (Figure 3, Panel 1A,B; *p* < 0.041). Detectable uPAR was also reduced at days 4 and 7 in SARS-CoV-2 MA30 infected C57Bl/6 mice treated with PEGSerp-1 (Figure 3, Panel 1C;*p*< 0.0215 and Panel 1D;*p* < 0.001, respectively). Comparison of alveolar and bronchial staining was equivalent for the inhibitory effects of PEGSerp-1 treatment on detected levels of cellular uPAR (Figure 3, Panel 1 D; *p* < 0.001). The hemorrhagic areas of lung consolidation in control virus-infected mice were also heavily stained for uPAR and this was similarly reduced by PEGSerp-1 treatment as might be expected given the fact that PEGSerp-1 binds uPA as a competitive inhibitor. At days 2 and 4 follow up in MA10 and MA30 SARS-CoV-2 infected mice, respectively, there was also a significant reduction in C5b-9 membrane attack complex (MAC) staining in infected mice receiving PEGSerp-1 treatment (Figure 3, Panel 1E-G; *p* < 0.0165 day 2 and *p*< 0.0056 for day 4 follow up). At day 7 in MA30-infected mice, however, the reduction in C5b/9 MAC staining with PEGSerp-1 treatment in MA30-infected mice was no longer significant (Figure 3, Panel 1H; *p* = 0.1201)

**Figure 3.**
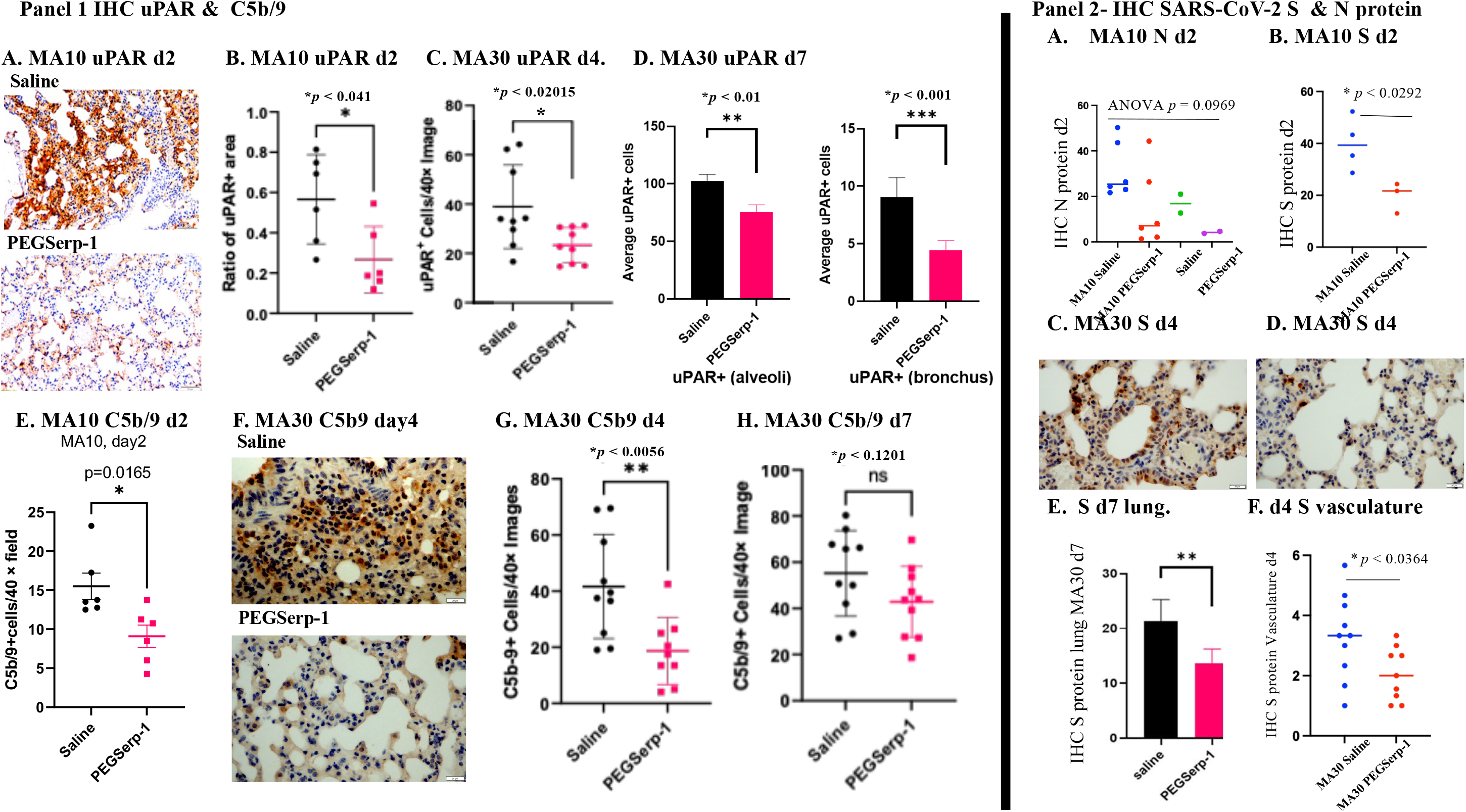
IHC stained sections demonstrate reduced uPAR, C5b-9 and SARS-CoV-2 S protein staining with PEGSerp-1 treatment in MA10 infected BALB/c and MA30 infected C57BL/6 mouse models. ***Panel 1*** - PEGSerp-1 treated mouse lung sections from day 2 follow up have significantly reduced detected uPAR positive stained areas (A) with reduced positively stained hemorrhagic areas in lungs at 2 days in MA10 infected BALB/c (B; *p* < 0.041) and reduced uPAR + stained cell counts at 4 days (C; *p* < 0.02015) and 7 days (D in alveoli (*p* < 0.01) and bronchi *(p* < 0.001) in MA30 infected C57BL/6 mice. C5b-9 MAC positive staining is reduced with PEGSerp-1 treatment in MA10 infected BALB/c mice (E; *p* < 0.0165) with PEGSerp-1 treatment. C5b-9 staining is reduced in MA30 infected C57BL/6 mice on IHC stained sections (F) with significant reductions at day 4 (G; p <0.0056) but a nonsignificant trend at day 7 follow up (H; *p* = 0.120). ***Panel 2*** – Reduced S protein detection with PEGSerp-1 treated MA10 infected BALB/c mice and MA30 infected C57BL/6 mice SARS-Cov-2 N protein is increased in MA10 infected mice with a nonsignificant trend toward a reduction with PEGSerp-1 treatment (A; *p* = 0.9069), whereas PEGSerp-1 treatments associated with reduced detectable S protein on IHC staining (B; *p* < 0.0292). In MA30 infected C57BL/6 mice increased S protein seen on IHC staining in lung sections from Saline treated mice (C) is reduced with PEG@erp-1 treatment at 4 days follow up)C, D. S IHC detection is reduced on micrographs of lung sections with PEGSerp-1 treatment (D). Cell counts positive for SARS-CoV-2 S protein staining in lung at 7 days (E; *p* < 0.01) and vascular sections at day 4 (F; *p* < 0.0364) with PEGSerp-1 treatment

uPAR and C5b-9 staining were similarly assessed in the myocardium and vascular tissues in SARS-CoV-2 infected mice. There is a nonsignificant trend toward a reduction in detectable uPAR staining at day 4 and day 7 follow up in the myocardium (Supplemental Figure 1, Panel 2I,J; *p* = 0.2675 and *p* = 0.2002, respectively). There is a trend toward a decrease in uPAR in the vasculature (Supplemental Figure 1, Panel2K, L; *p* = 0.0867), however uPAR detection was significantly reduced in the myocardial vessels at day 7 (Supplemental Figure 1, Panel 2L; *p* < 0.0082). Detectable C5b-9 staining was not significantly reduced at days 4 and 7 in the myocardium and vasculature of PEGSerp-1 treated virus-infected mouse heart tissues (*p* = 0.1525 and *p* = 0.2725, respectively). This suggests that changes in the myocardium and associated vessels, if present, occur later than in the lungs as might be predicted based upon the known route of SARS-CoV-2 infection.

### Immunohistochemical staining for viral N and S proteins in mice infected with MA10 or MA30

The presence of SARS-CoV-2 spike (S) and nucleocapsid (N) proteins were also assessed by IHC analysis of lung tissues in infected mice. As expected, detectable S and N protein levels on IHC stained sections were increased in virus-infected mice when compared to uninfected control mice (Figure 3, Panel 2A). N protein levels were not reduced with PEGSerp-1 treatment in MA10-infected mice at 2 days follow up, although there was a trend toward a reduction (Figure 3, Panel 2A; *p* = 0.0969). Significant reductions in staining for S proteins were detected at day 2 in MA10-infected mice (Figure 3, Panel 2B; *p* <0.041) and day 7 in MA30 infected mice in lung (Figure 3, Panel 2C-E; *p* < 0.001 at day 7) with PEGSerp-1 treatments. No detectable reduction in S protein was seen on day 4 *(p* = 0.5105), but detectable S protein staining was reduced in lung samples in MA30 infected mice with SARS-CoV-2 staining (Figure 3, Panel 2E; *p* < 0.001). Detectable S protein staining was not reduced in the myocardium of SARS-CoV-2 MA30 infected mice (*p* = 0.8666), however, detected S was significantly reduced in the vasculature on IHC at 4 days follow up (Figure 3, Panel 2F; *p* < 0.0364).

### Analysis of gene expression changes in the lung by qRT-PCR analysis in SARS-CoV2 infections

To further assess the level of virus in the lungs following IN infection, SARS-CoV-2 E and N gene expression in frozen mouse lung tissue at day 4 in MA30 infected mice were measured by qRT-PCR array. Increased expression of E (Figure 4, Panel 1A; *p* < 0.0017) and N gene expression (Supplemental Figure 4R,S) were detected in all SARS-CoV-2 MA30 infected mice at 4 days follow up, when compared to uninfected lung isolates. There was a corresponding significant increase in the canonical Interferon (IFN)-upregulated cellular gene ISG15 in MA30 infected mice compared to uninfected lung tissues, (Figure 4, Panel 1B; *p* < 0.0044). However, PEGSerp-1 treatment did not alter detectable levels of SARS-CoV-2 E gene (Figure 4, Panel 1A; *p* = 0.9217 and Supplemental Figure 4R, S; *p* = 0.4138 and 0.4847), nor the virus-induced upregulation of ISG15 gene expression at 4 days follow up (Figure 4, Panel 1B; *p* = 0.9260). This suggests that PEGSerp-1 is not directly impacting the virus, but instead targeting, as expected, the inflammatory response to infection.

**Figure 4.**
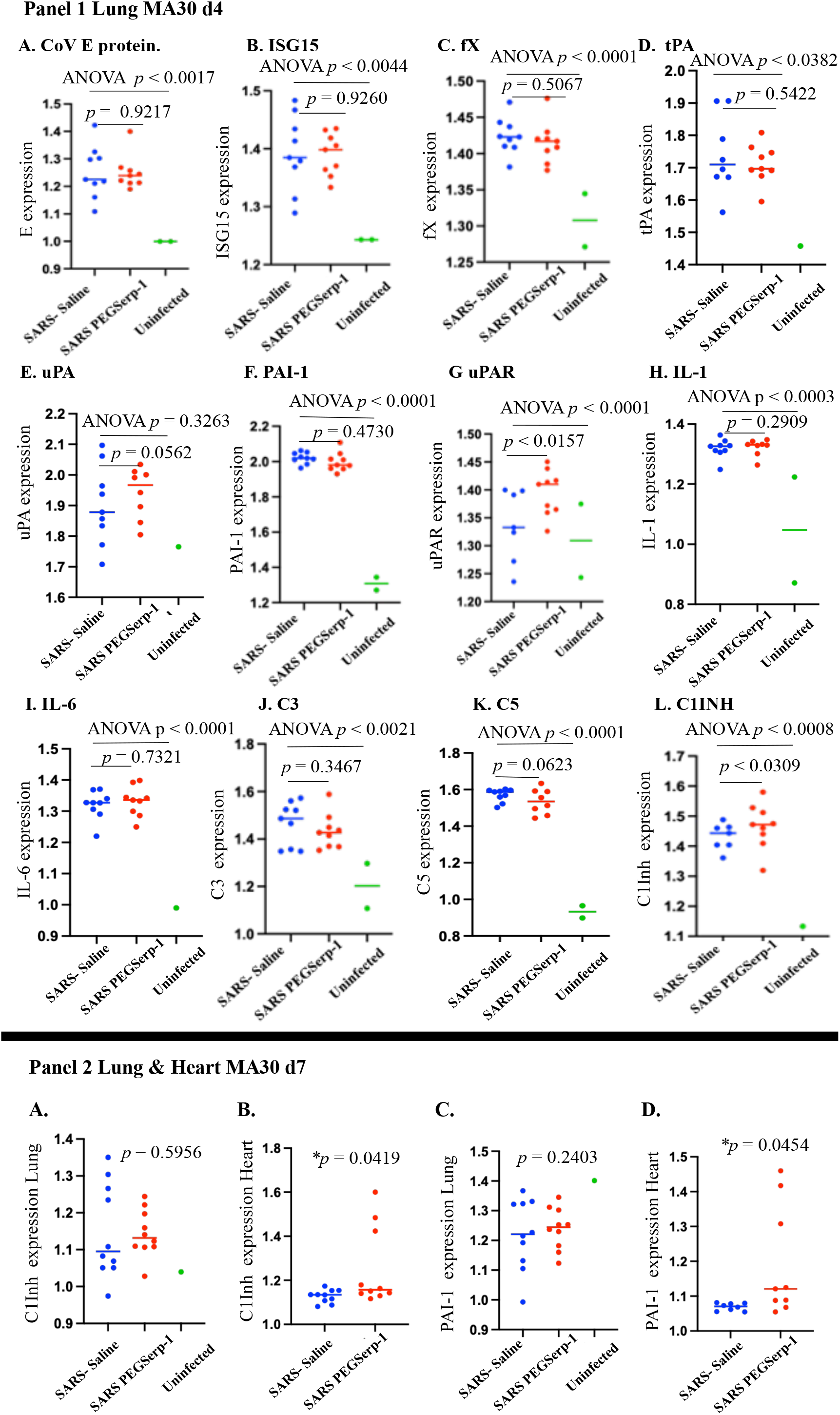
Relative gene expression for SARS-CoV-2 and associated inflammation are increased after SARS-CoV-2 MA30 infection in lung and heart tissues PEGSerp-1 treatment significantly modified gene expression for uPAR and complement pathways. ***Panel 1*** - Relative gene expression of MA30 E (A; *p* < 0.0017)) and ISG15 (B; p < 0.0044)) are significantly increased in infected lung extracts at 4 days follow up in MA30 infected mice, when compared to uninfected mouse lungs. Factors in the coagulation pathway, fX (C; *p* < 0.0001), tPA (D; *p* < 0.0382), PAI-1 (F; *p* < 0.0001) and uPAR (G; p < 0.0001) were increased in infected mouse lung extracts when compared to uninfected lungs with SARS-CoV-2 MA30 infection. uPA was not increased with infection (E;*p* < 0.3363). IL-1 (H;*p* < 0.0003), IL-6 (I;*p* < 0.0001), C3 (J;*p* < 0.0021), C5 (K;*p* < 0.001) and C1INh (L; *p* < 0.0008) were also significantly increased with infection. PEG Serp-1 treatment produced a borderline decrease in C5 (K; p= 0.0623) and a significant increase in uPAR (G; p < 0.0157) and C1Inh (L; p < 0.0309). ***Panel 2*** - At 7 days follow up, C1Inh relative gene expression is no long increased with PEGSerp-1 treatment in MA30 infected mouse lungs at 7 days (A) but was significantly increased in heart extracts with PEGSerp-1 treatment at 7 days (B; p < 0.0419). PAI-1 gene expression was not increased in lung extracts with PEGSerp-1 treatment at 7 days follow up in MA30 infected mouse lungs (C; *p* = 0.2403), but PAI-1 expression is significantly increased at 7 days follow up in heart extracts (D; *p* < 0.0454).

To assess gene expression in multiple acute response coagulation and inflammatory response pathways qRT-PCR array analysis was performed on SARS-CoV-2 MA30 infected, in parallel with uninfected lung tissues. Increased gene expression was observed with a general increased at 4 days with infection when compared to 2 days and with a subsidence by day 7 (Supplemental Figure 2A). Specifically, in the coagulation pathways; Significant increases were seen for factor X (fX; Figure 4, Panel 1C;*p* < 0.0001), tPA (Panel 1D;*p* < 0.0382), PAI-1 (Panel 1F;*p* < 0.0001), and uPAR (Figure 4, Panel 1G; *p* < 0.0029) gene expression, but not for either uPA (Figure 4, Panel 1E; *p* = 0.3263) or neuroserpin *(p* = 0.1883) in the lungs of virus-infected mice. Gene expression for inflammatory cell response pathways were also significantly elevated for tumor necrosis factor (TNF; Supplemental Figure 2C; *p* < 0.0044), Interleukin-1 (IL-1; Figure 4, Panel 1H; *p* < 0.0003), IL-6 (Figure 4, Panel 1I; *p*< 0.0001), IL-10 (Supplemental Figure 2D; *p* < 0.0001), complement C3 (Figure 4, Panel 1J; *p* < 0.0021), C5 (Figure 4, Panel 1K; *p* < 0.0001), C1INH (Figure 4, Panel 1L, *p* < 0.0008) and vascular endothelial growth factor (VEGF; Supplemental Figure 2E, p < 0.0222), but not for Transforming growth factor (TGF-beta; Supplemental Figure 2D; *p* = 0.4695) in infected mouse lung tissue when compared to uninfected mice.

A detectable change in gene expression in select coagulation and complement pathway inflammatory markers was seen with PEGSerp-1 treatment in virus-infected mice. PEGSerp-1 significantly increased uPAR (Figure 4, Panel 1G, *p* < 0.0157), produced a borderline decrease in C5 (Figure 4, Panel 1H; *p* < 0.0623), and significantly increased C1Inh (Figure 4, Panel 1L; *p* < 0.0309). Nonsignificant inhibitory trends were observed for TNF and C3 with PEGSerp-1 treatments (Supplemental Figure 4B,O). The increased uPAR and the reductions in C3 and C5 and increases in C1Inh support a greater effect of PEGSerp-1 treatment on the uPAR and complement pathways in the lungs of virus-infected mice.

Formalin fixed paraffin embedded (FFPE) lung samples from the SARS-CoV2 MA10 (48 hours) and the SARS-coV-2 MA30 (7 day) were also assessed for gene expression changes with SARS-CoV-2 MA10 (48 hours) and SARS-CoV-2 MA30 (Day 7) infections. Heart samples were additionally available for the day 7 SARS-CoV-2 MA30 samples. The MA10 SARS-CoV-2 MA10 infection lung samples, 48 hours, were limited and sample numbers too small to provide statistically meaningful findings in the MA10 infection samples. However, there is an apparent association between inflammatory markers and time of follow up with greater increases overall at day 4 over day 2 and then a subsidence of levels by day 7 for several markers, consistent with the overall changes in clinical score and weight loss with SARS-CoV-2 infections (Supplemental Figure 2A). For the day 7 FFPE samples we saw similar trends in gene expression as seen in the day 4 samples with PEGSerp-1 treatments in MA30 infected mice. C5 was again reduced with PEGSerp-1 treatments at day 7, with a nonsignificant trend (Supplemental Figure 2M; *p* < 0.1601), whereas there was a borderline significant change in day 4 follow up samples (Figure 4, Panel1K; *p* = 0.0623). uPAR which was significantly increased at 4 days with PEG Serp-1 treatment was no longer increased at 7 days follow up (Supplemental Figure 4G; *p* = 0.1514). C1Inh, which was significantly increased with PEGSerp-1 treatments at 4 days in the lungs, was no longer significantly increased at 7 days in lung sections (Figure 4, Panel 2A; *p* = 0.5956).

Heart tissues in contrast had a significant increase in day 7 cardiac samples for inflammatory markers after MA30 infection while lung samples were showing a reduction (Supplemental Figure 4). In the cardiac samples, and of interest for serpin treatments, in the day 7 follow up heart tissue samples C1Inh was significantly increased on qPCR array analysis with PEGSerp-1 treatments (Figure 4, Panel 2B; *p* < 0.0419). Similarly, the PAI-1 mammalian serpin, while no longer increased in lung samples (Figure 4, Panel 2C; *p* = 0.2403) was now increased in cardiac samples at day 7 follow up (Figure 4, Panel 2D; *p* < 0.0454). uPAR and C5 were not significantly altered in the heart sections on day 7 (Supplemental Figure 4H, N; *p* = 0.2184 and *p* = 0.1617, respectively). This suggests that PEGSerp-1 treatment modified gene expression of uPAR and complement pathways with associated changes in regulatory serpins in both lung and heart tissues with differing activation time windows.

## 4. Discussion

With this study we examine PEGSerp-1 treatment of severe immune-mediated lung damage in SARS-CoV-2 viral infections. We demonstrate a significant improvement in clinical scores and weight gain during systemic treatment with PEGSerp-1, a Myxomavirus derived serpin, in two different mouse-adapted models of SARS-CoV-2. This unique serpin targets activated serine proteases in both coagulation and complement pathways, significantly improving weight and clinical score with reduced immune cell invasion in the lung and heart. SARS-CoV-2 infections begin in the nasopharynx and quickly migrate to the lungs with subsequent invasion of other visceral organs with a damaging immuno-thrombotic reaction. Parallel changes in both heart and lungs of PEGSerp-1 treated mice indicate a systemic and sequential reduction in immune responses to SARS-CoV-2 infection. Inflammatory M1 macrophage infiltrates were significantly reduced with PEGSerp-1 treatments during infection in both MA10 infected BALB/c mice and in MA30 infected C57Bl/6 mice. M2-activated macrophages as well as CD4 and CD8 T cell responses were modified later during MA30 infections. C57BL/6 and BALB/c mice have marked differences in their immune response patterns to virus infections (*ie*. C57-derived mice tend to be Th1-polarized whereas BALB/c-derived mice are more Th2-polarized) (54). Thus, PEGSerp-1 suppressed SARS-CoV-2 virus-induced pathologic immune reactions in multiple genetic backgrounds.

These findings are consistent with prior reported improved lung inflammation, vasculitis and associated improved survival in Serp-1 treated MHV68 gamma-68 herpesvirus-infected mouse model (42,43). PEGSerp-1 treatment thus has the potential to improve outcomes in both RNA and DNA viral lung infections. Balancing activation of the thrombotic/thrombolytic and complement pathways via serpin treatment has the potential to restore homeostasis in severe viral infections where there is aggressive and damaging activation of immune and thrombotic proteases. Key serine proteases in each of these molecular pathways are regulated by endogenously expressed mammalian serpins (44). Serpins are suicide inhibitors that target activated serine proteases and act as enzyme pseudo-substrates at sites of immune and coagulation pathway activation. These protease pathways are routinely activated during severe virus infections (36–40). Two mammalian serpins, alpha 1 antitrypsin (A1AT, SERPINA1) and Complement 1 Inhibitor (C1INh, SERPING1) are in clinical trials as potential treatments for SARS-CoV-2 ARDS (39,40). Anti-thrombin III (ATIII, SERPINC1) was previously investigated as a treatment for bacterial sepsis and associated disseminated intravascular coagulation (DIC). ATIII treatment exhibited a trend toward improved outcomes in bacterial sepsis, but did not result in statistical significance, whereas C1Inh treatment has early benefit (38,40).

PEGSerp1 is a myxoma poxvirus-derived serpin with high efficacy that has developed through millions of years of virus evolution to protect the virus from activated inflammatory cellular attack by host macrophages and monocytes (41–45, 47–50). Serp-1 targets multiple immune and thrombotic pathways that are activated in severe viral lung infections and septic states with the potential to restore homeostatic balance in excessively activated uPAR and complement pathways. Inflammatory cell invasion and increased pro-inflammatory gene expression were both detected in MA virus-infected mouse lung tissue when compared to normal uninfected lung tissues. PCR array analyses demonstrated a clear and significant increase in viral protein genes, as well as interferon mediators such as ISG15 and a range of immune response thrombotic and immune response pathway members in the lungs of MA SARS-CoV-2 infected mice. However, both IHC and qRT-PCR analyses demonstrated that PEGSerp-1 targeted uPAR in the thrombolytic pathway, and C5 and C1Inh in the complement pathway were significantly altered.

Mechanistically, Serp-1 anti-inflammatory activity is dependent on the uPA receptor (uPAR) (47,48) and Serp-1 loses all therapeutic effects in uPAR-knockout mice (48). The uPA/uPAR complex activates plasmin and matrix metalloproteinases (MMPs), acting both as a thrombolytic protease and an immune modulator that regulates macrophage and neutrophil activation and tissue invasion. Of interest, the effects of PEGSerp-1 treatment in the lungs of MA SARS-Cov-2 infected mice were more diverse than expected: uPAR and C1Inh both increased, while C5 was borderline reduced. In the heart, PAI-1, a uPA regulating mammalian serpin was increased with PEGSerp-1 treatment at day 7 follow up. In prior work with the MHV-68 gamma herpesvirus infection model, there was a significant increase in serpins that target the uPA pathway (43). Serp-1 binding to uPAR on the surface of activated and mobilized macrophages was previously demonstrated and may further alter various downstream cellular activation and gene expression through interactions with proteins associated with the uPAR lipid rafts (47). The PEGSerp-1 induced changes in complement gene expression may be more consistent with reduced levels of lung inflammation and tissue damage, however these analyses were performed on whole lung isolates at a single time after infection and may or may not represent changes in protein expression and activity in either infiltrating immune cells or resident lung cells. Thus, these detectable protective changes induced by PEGSerp-1 are a composite of multiple tissues inclusive of lung epithelium, arterial endothelium and invasive immune cells.

The SARS-CoV-2 spike (S) protein binds to cellular ACE2 receptors and during entry at the cell surface is cleaved at the S2’ site by the surface transmembrane protease serine 2, TMPRS2, which promotes fusion of the viral envelope with the cell membrane (9,32–35). The S protein has been reported to be cleaved by circulating thrombotic proteases, thrombin and factor X, as well as the thrombolytic protease plasmin (31–36). A reduction in S protein was observed on IHC stained sections in PEGSerp-1 treated MA infected mice. Our qRT-PCR analyses demonstrated a clear increase in SARS-CoV-2 E and N gene expression in the lungs of SARS-CoV-2 MA30 infected C57Bl6 mice, but no change was seen in PEGSerp-1 treated animals. The reduced detectable S protein with IHC staining is likely an indirect result of the serpin-mediated amelioration of lung damage and inflammation. Further analyses will be required to assess the potential for PEGSerp-1 to alter SARS-CoV-2 S protein cleavage and any impact on infection, thus we postulate that PEGSerp-1 therapeutic effects are primarily anti-inflammatory and not anti-viral.

Severe SARS-CoV-2 viral lung infection is associated with ARDS, especially when patients reach the ICU. Immune mediated damage and widespread micro-thrombotic occlusions cause severe lung and multiorgan damage. With the most severe respiratory or mosquito-borne viral pandemics, including from dengue, influenza, Ebola and SARS-CoV-2, high mortality rates result until effective vaccines and therapeutic treatments are developed. Treatments for the early viremic phase of COVID-19 have advanced with monoclonal antibodies and anti-viral therapeutics, but there is still a need for more effective immune modulators during the later stage pro-inflammatory phase of the disease (1–8, 11–41). Drugs targeting pro-inflammatory cytokines such as IL-1 and IL-6, as well as JAK pathway inhibitors, have been effective, but mortality for high-risk patients remains high. Dysregulation of both the thrombotic and thrombolytic pathways co-activate excessive innate immune responses and complement activation cascades which in turn further damages cells and increases systemic thrombosis. Activation of uPAR (especially as assessed by the upregulation of circulating cleaved suPAR) and the complement cascade are markers for progression to severe COVID-19 complications. Serine proteases regulate these innate immune response pathways and, in addition, the cell surface serine protease, TMPRSS2. Mammalian serpins represent 2-10% of circulating blood proteins, including antithrombin III (AT), alpha-1 antitrypsin (AAT), and complement inhibitor (C1Inh), all of which regulate thrombosis and inflammation. Highly potent serpin-based immune regulators have also been evolved over millions of years in viruses and bacteria by selectively targeting sites of host serine protease activation that normally help combat pathogens during infection (44,45,53–55). Thus, the poxvirus-derived Serp-1 protein evolved in myxoma virus as a secreted inhibitor of activated myeloid cells. Indeed, activated macrophages more effectively recognize and clear myxoma virus from infected tissues when the viral Serp-1 gene is deleted (53,54).

In this study, we have examined treatment of two mouse adapted SARS-CoV-2 models using PEGSerp-1, a pegylated version of the viral Serp-1 serpin that targets multiple serine proteases in the coagulation and complement protease cascades. Treatment significantly improved clinical scores, decreased lung inflammation and consolidation and modified uPAR and complement levels. In previous work, we showed that systemic treatment with purified virus-derived Serp-1 protein improves survival and significantly reduces inflammation in lethal mouse models infected with gamma herpesvirus (MHV68) or Ebola (43). Additionally, in a pristane-induced lupus alveolar hemorrhage model, Serp-1 treatment also improved pathologic lung hemorrhage and reduced macrophage invasion (49). Purified Serp-1 protein expressed in CHO cells has also been tested and proven safe and effective in a randomized Phase IIa clinical trial for treating cardiovascular patients with unstable coronary disease and stent implant where markers for heart damage, troponin T and creatinine kinase MB were significantly reduced (MACE of zero) (50).

Overall, the study described here for SARS-CoV-2 and earlier studies with other models for severe viral infection strongly support further studies and development of PEGSerp-1 as a new antiinflammatory therapeutic or biologic for advanced virus-induced lung and vascular damage, designed to target coagulation disorders, micro-thrombi, hemorrhage and damaging inflammation. Longer term studies are needed to assess the impact of PEGSerp-1 treatment on severe viral ARDS, immune-thrombotic complications and chronic infections that may contribute to chronic debilitating disease, as seen in long COVID following SARS-CoV-2 infection.

## 5. Methods

### SARS-CoV-2 Infection

All animal studies conformed to local and national guidelines for animal care and experimentation and were approved by the Institutional Animal Care and Use Committee (IACUC) of Arizona State University (ASU) (#20-1761R). Animals were housed in barrier conditions at the ASU Animal Care Services vivarium and bred under specific pathogen-free conditions. Equal numbers of male and female mice were used for all studies. Mice were weaned at 3 weeks, maintained on a 12-hour light-dark cycle and fed water and standard rodent chow *ad libitum*. Mouse cohorts are transferred from the ABSL1 colony to a separate ABSL2 colony until beginning work in the ABL3 facility. C57BL/6 mice were purchased from the Jackson Laboratory and BALB/c mice were bred in house. Equal numbers of male and female mice were used for the studies. Mice were infected at 8-10 weeks age and treated daily with daily intraperitoneal injection (IP) of either Saline control or PEGSerp-1 100ng/gm body weight with treatment continued throughout the course of infection to follow up. For each study either BSL3 treatments (YL, KK) or the histopathology were blinded to treatment. For euthanasia, mice were deeply anaesthetized with an overdose of 80 mg sodium pentobarbital/kg body weight. Lung and heart were harvested after euthanizing. Tissues for pathology analyses were post-fixed in formalin and tissues for qPCR were frozen for trizol.

### Virus expression, purification and titer titers

i. For SARS-CoV-2 MA10 infection - Mouse adapted SARS-CoV-2 MA10 was obtained from Dr Ralph Baric’s lab (University of North Carolina at Chapel Hill) and was propagated in Vero TMPRSS2 cells (gift of Stefan Pöhlmann)(48). For MA10 infection 2×10^5^ pfu was inoculated by intranasal route (IN) in BALB/c mice (51).
ii. For SARS-CoV-2 MA30 infection - Mouse adapted (MA) SARS-CoV-2 (LD 50 - 1.3 X 10^3^ pfu) was inoculated IN in C57Bl/6 mice as previously described (52,53). For IN instillation, mice were sedated prior to infection with isofluorane and infected IN with virus (1.3 X 10^3^ pfu) in 5μl DMEM, as previously described (52,53). Mice were monitored until stable after infection and checked twice daily. Mice are kept warm with warm blankets if recovery is prolonged. Animals are monitored for activity, posturing, respiratory distress, decreased appetite and weight loss on comparison to control animals as warning markers for potential need to euthanize the test animals earlier than the projected dates. Mice are weighed daily. Mouse-adapted SARS-CoV-2 SARS2-N501Y_MA30_ was propagated in A549-huACE2 cells (51).

Human A549 cells (Verified by ATCC) were cultured in RPMI 1640 (Gibco catalog No. 11875) supplemented with 10% FBS, 100 U/mL of penicillin, and 100 μg/mL streptomycin. The generation of A549-ACE2 cells was described previously (53).

### PEGSerp-1 - PEGylation and protein purification

Serp-1 (m008.1L; NCBI Gene ID# 932146) was expressed in a Chinese hamster ovary (CHO) cell line (Lucas lab)(43,47–50). Serp-1 protein used in this research is GMP-compliant >95% pure, as determined by Coomassie stained SDS-PAGE and reverse-phase HPLC; endotoxin-free by LAL (limulus amebocyte lysate) assay.(Supplemental Figure 3) For PEGylation, Serp-1 is incubated with mPEG-NHS (5 K) (Nanocs Inc., #PG1-SC-5k-1, NY) in PBS buffer (pH 7.8) at 4°C overnight according to standard PEGylation protocols (49). PEGSerp-1 is purified by FPLC using an ÄKTA pure protein purification system with Superdex-200.

### Histological analysis

Tissues were analyzed by routine hematoxylin and eosin (H&E) staining as well as immunohistochemical (IHC) staining. Total inflammatory cell infiltrate area, hemorrhage when present and consolidation areas were measured and normalized to the total lung area examined (43, 47–49). Three high power fields were assessed for each specimen and the mean for positively stained cell counts per 40X field calculated. IHC was performed for the detection of immune cell markers as well as uPAR and C5b-9 MAC: F4/80 for macrophage; CD3 for nonspecific T lymphocytes; Ly6G for neutrophils; CD19 for B lymphocytes and C5b/9 antibody (Abcam, ab 55811), and uPAR (R&D Systems, AF534,1:100). For IHC, the primary antibodies included the following: Anti-mouse C4d Cat: HP8033; Ra pAb to F4/80 ab100790; Rb pAb to CD3m ab 5690 and secondary antibody: Goat antiRat IgG2a Hrp conjugated Cat: A110-109P. Sections were examined using an Olympus BX51 microscope with 4x–100x objectives, a Prior ProScan II stage and Olympus DP74 CMOS camera and cellSens software analysis system. Staining for C4d was used as a secondary marker of rejection.

An independent pathology score was also performed blinded (SG), with score is based on intra-alveolar edema and chronic inflammatory cells (lymphocytes, macrophages). If the airspaces were clean and open without significant inflammation or edema, Score assigned was 1+. For airspaces partially open, with increased chronic inflammation with or without edema, Score assigned 2+. For sections with a majority of airspaces consolidated, with marked chronic inflammation with or without edema, Score assigned 3+.

### RNA isolation and qPCR

Samples were assayed for gene expression in infected lung and heart tissues, with and without PEGSerp-1 treatment as well as in normal uninfected lung and heart isolates. Lung and heart samples from controls or SARS-CoV-2 MA30-infected mice were collected four days post infection and frozen. Samples were also collected from formalin fixed, paraffin embedded lung and heart samples at 2 and 7 days follow up. For RNA extraction, tissues were harvested in RNA STAT-60 (Tel-Test) and disrupted by bead homogenization (Omni, INC). Isolated mouse lungs were transferred to Trizol (Thermo Fisher Scientific). RNA was extracted using the Qiagen RNAeasy Mini Kit (Qiagen, 74106), for frozen tissues and using the Qiagen RNAeasy kit (Qiagen 73504) for FFPE specimens, according to the manufacturer’s instructions. Briefly, 0.2 mL chloroform was added to 1 mL of RNA STAT-60, samples were mixed for 30 seconds, kept on ice for 5 min and centrifuged 12,000g for 15 minutes at 4 °C. 0.5 mL of 100% ethanol was added to aqueous phase (RNA), samples were kept for 5 min at 4 °C and centrifuged as above. The final RNA pellet was resuspended in molecular biology grade water (Sigma). Total RNA was reverse transcribed with an oligo (dT) primer and a Moloney murine leukemia virus reverse transcriptase (MMLV RT, TakaraBio). cDNA was analyzed by quantitative PCR amplification using SYBR Green qPCR Master Mix (QuantaBio) on a Bio-Rad CFX96 Real-Time PCR Detection System. Primers were designed to amplify mRNA-specific sequences, and analysis of the melt-curve confirmed the amplification of single products. Uninfected samples were used as controls. Relative expression was normalized to GAPDH. Primer sequences used are provided in Supplemental Table 2. Primers were designed using the IDT website, PrimerQuest Tool, for qPCR and confirmed in the literature.

Primers used for qPCR included SARSCoV-2 E and N genes, Gapdh, Isg15, Fx, Tpa, Upa, Mmp2, Mmp-9, Upar, Pai-1, Nsp, Tnf, Il-1, Il-6, Il-10, Tgfb, Vegf, C3, C5, C1inh SARS-Cov-2 E and N gene primers were provided by Dr B Hogue and Gapdh standard by Dr M Rahman. Real time qPCR was analyzed on a BioradCFX96 Real Time PCR array with C1000 Touch Thermal cycler.

### Statistical Analysis

Immunohistochemical analyses were initially read blinded to treatments and mouse strain. In the initial MA30 studies, the histology was evaluated by investigators blinded to treatments. In the second MA30 studies both investigators performing the mouse studies as well as initial histological analyses were performed blinded. Significance in each parameter was assessed by StatView version 5.0.1 (SAS Institute, Inc.) or Graph Pad Prism using analysis of variance (ANOVA) with Fischer’s LSD (Least Significant Difference) comparison, or a Student’s unpaired *t*-test *(p* < 0.05 was considered significant). If more than one analysis was performed, scores and measurements were combined. The areas with greatestpositive staining were used for IHC cell counts. The mean for three areas of consolidation and the mean for three positively stained cell counts in lung or heart microscopy sections at 40X high power field were calculated and used for statistical analysis.

## 6. Acknowledgements

**We would like to acknowledge the following funing agencies** - BGH - Tohono O’Odham Nation Seed Award GR40818; GM, MR, BGH - Mercatus Center Fast Grant (G09202-300); AL - NIH R01AR074627-01A1 and ASU Skysong Innovations. We would like to thank Dr. R. Baric for providing the SARS-CoV-2 M10 and Dr. S. Perlman for the SARS-CoV-2 MA30 virus constructs.

