## Supplementary figures and images for "Treatment with anti-inflammatory viral serpin modulates immuno-thrombotic responses and improves outcomes in SARS-CoV-2 infected mice"

### Supplemental Figure 1

Supplemental Figure 1. Immune cell responses

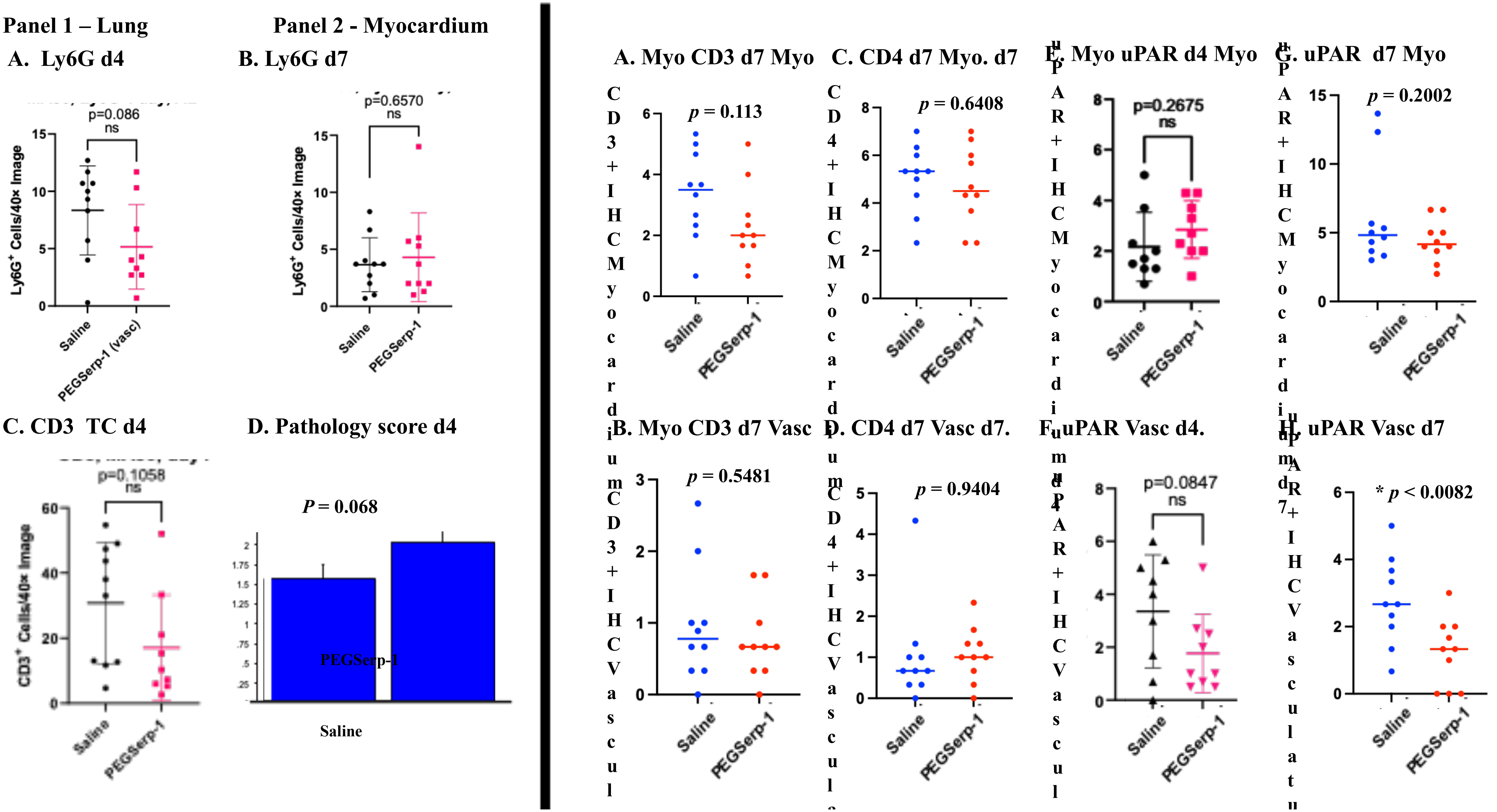

### Supplemental Figure 2

#### Supplemental Figure 2 - qPCR

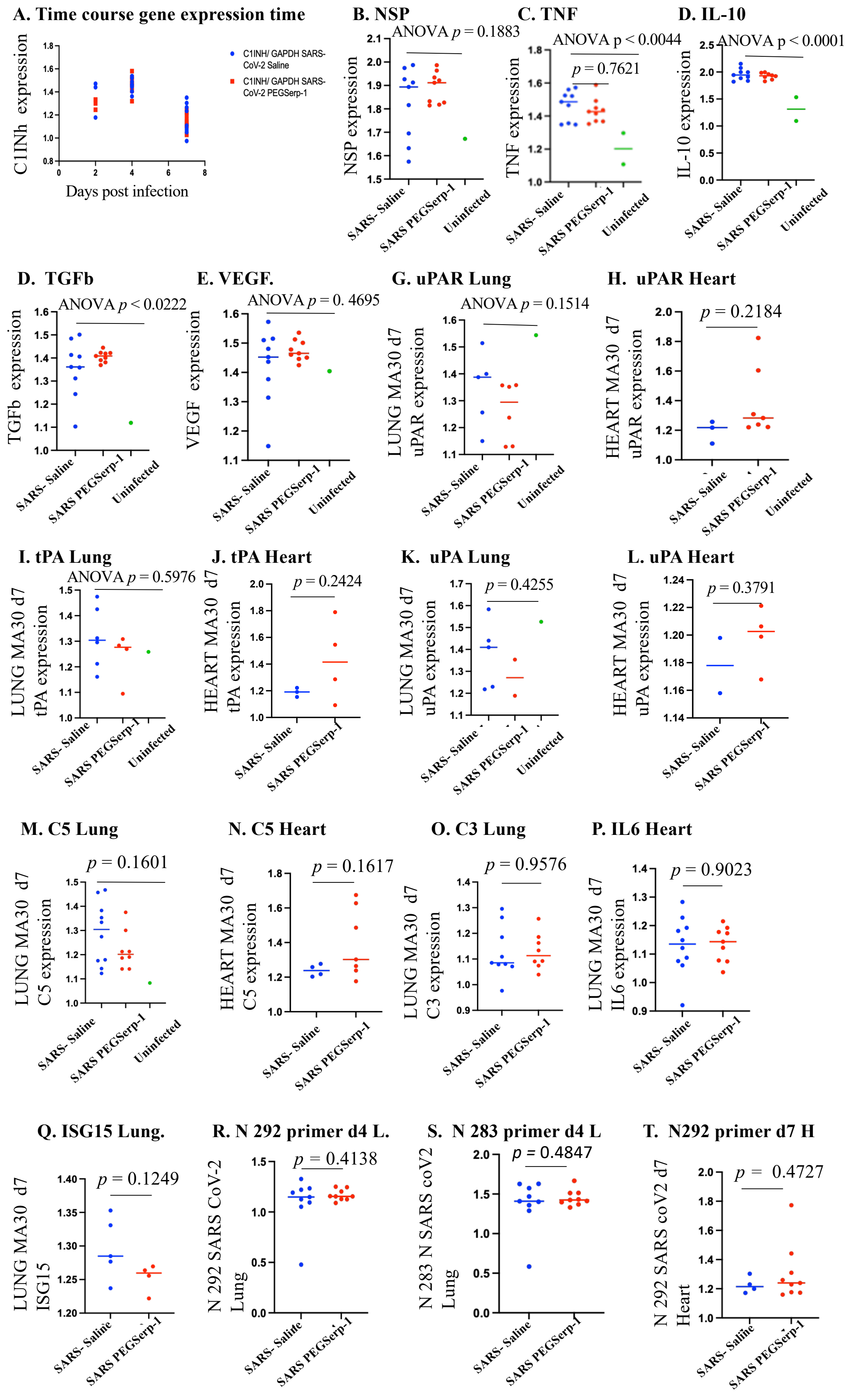

### Supplemental Figure 3

**Supplemental Figure 3. PEG Serp-1 binding to uPA**

**Western blot**

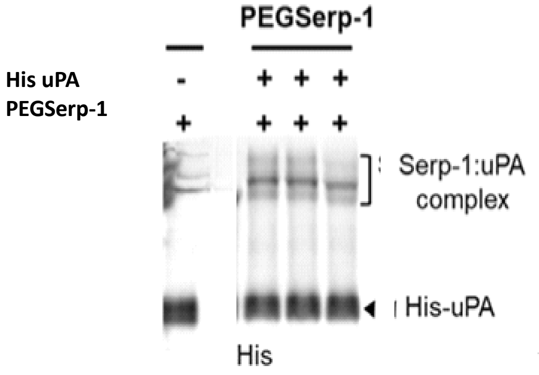
