## Supplemental Table 1 for "Treatment with anti-inflammatory viral serpin modulates immuno-thrombotic responses and improves outcomes in SARS-CoV-2 infected mice"

**Table 1 MS analysis - PEGSerp-1 treated mouse DAH lung tissue**

| Accession | Description | Symbol | # Peptides |
|---|---|---|---|
| P01027 | Complement C3 | C3 PE | 34 |
| B7FAV1 | Filamin, alpha | Flna PE | 23 |
| P60710 | Actin, cytoplasmic 1 | Actb PE | 17 |
| A0A0R4J0X5 | Alpha-1-antitrypsin 1-3 | Serpina1c PE | 13 |
| E9PV24 | Fibrinogen alpha chain | Fga PE | 12 |
| P20918 | Plasminogen | Plg PE | 11 |
| P01029 | Complement C4-B | C4b PE | 9 |
| P98086 | Complement C1q subunit A | C1qa PE | 4 |
| P04186 | Complement factor B | Cfb PE | 4 |
| Q8CG14 | Complement C1s-A | C1sa PE | 4 |
| P14106 | Complement C1q subunit B | C1qb PE | 3 |
| Q8CG16 | Complement C1r-A | C1ra PE | 3 |
| Q02105 | Complement C1q subunit C | C1qc PE | 2 |

*Average Xcorr >4.0
